# Experimental evolution of *Vibrio cholerae*: Identification of genes involved in motility in presence of polymyxin B

**DOI:** 10.1101/2021.07.12.451780

**Authors:** Sean Giacomucci, Antony T. Vincent, Marylise Duperthuy

## Abstract

*Vibrio cholerae* is the bacteria responsible for the cholera disease and a natural inhabitant of aquatic environments. Biofilm formation is important for human colonization and environmental survival. Motility is essential for adhesion and biofilm formation by *V. cholerae*. In a previous study, we showed that motility and biofilm formation are altered in the presence of sub-inhibitory concentrations of polymyxin B in *V. cholerae*. In this study, we performed an experimental evolution to identify genes rescuing the motility in the presence of polymyxin B. Mutations in 5 genes have been identified: *ihfA, vacJ* (*mlaA*), *mlaF, dacB* and *ccmH*. The details of these mutations, their potential impact on the function of the proteins they encode and on the motility in presence of polymyxin B are discussed.

## Introduction

*Vibrio cholerae* is a Gram-negative pathogenic bacterium responsible for the cholera disease, which infect millions of people per year worldwide. More than 200 serotypes have been reported but only the O1 and O139 cause the cholera [1]. The O1 strains are further divided in 2 biotypes: Classical and El Tor. The Classical biotype was responsible for 6 pandemics and has been replaced by the El Tor biotype in 1961, which is responsible for the ongoing 7^th^ pandemic [1]. The O139 serotype, which causes local epidemics, appeared in 1992 and is derived from the O1 El Tor biotype [2]. The non O1 / non O139 serogroups can cause less serious gastroenteritis [3].

For most of the pathogenic bacteria, motility is essential for host colonization and virulence [4]. Different types of motility exist: swimming, swarming, twitching, gliding and sliding, the latest being a passive process that does not involve energy [5]. Swarming, twitching, gliding and sliding are surface motility mechanisms whereas swimming allows bacteria to move in liquid suspensions and involves active propulsion by the flagella. Flagella are filamentous apparatus anchored to the bacterial envelope and divided in three major structures: the basal body, the hook and the filament [6]. The movement is ensured by the rotation of the flagella filament driven by a motor-like structure. In most of the bacteria, the motor uses a H+ gradient as energy source [7]. In *Vibrio*, the rotation of the flagella is driven by a Na+ motor [8], which provides a rotation speed up to 5 times faster than the H+ driven flagella of *E. coli* [9].

*V. cholerae* is motile and possess a single polar flagellum. The filament is composed of five homologous subunits: FlaA, FlaB, FlaC, FlaD and FlaE, and the flagellum is coated by an outer membrane sheet [10, 11]. Besides its crucial role in motility, *Vibrio* flagellum is also essential for biofilm formation, colonization and virulence [6]. In *V. cholerae*, the expression of the flagellar gene expression is tightly regulated and is organized in a 4-steps hierarchical process, resulting in 4 classes of genes. There is only one class I gene, the σ^54^-dependant master regulator of the flagella regulation hierarchy *flrA*. This regulator is essential for flagella synthesis and its expression is activated by the histone-like nucleoid structuring protein (H-NS or VicH) [12] and repressed by FlhG, which regulates the number of flagella [13]. FlrA control the expression of the class II genes that include genes encoding for structural components of the flagella such as the MS (membrane/supramembrane) ring, the ATPAse and the export apparatus. In addition, *flhG* and *flhF* also belongs to the class II genes and control the number and polar localization of the flagella [14]. Among the class II genes, *flrBC* and *fliA* are the main activators of the class III and class IV genes, respectively. The class III genes contain other genes encoding for proteins essential for the structure of the flagella including the inner membrane apparatus, the proximal and distal rod and the hook-related proteins, among others [6]. The FlaA flagellin gene (*flaA*) also belongs to the class II gene and is the only flagellin essential for motility [10]. The class IV genes contains those encoding for the other flagellins (*flaB, flaC, flaD* and *flaE*), the anti-sigma factor FlgM and the motor (MotA, MotB and MotX) [6]. FlgM is a negative regulator of *fliA* and prevent the expression of class IV genes until the flagella apparatus is successfully assembled and enable the secretion of FlgM [15].

In our previous study, we showed that the biofilm formation was impaired in presence of subinhibitory concentrations of polymyxin B, due to a significant reduction in the proportion of flagellated *V. cholerae* O1 and O139 [16]. In addition, a motility reduction was observed in the presence of polymyxin B and was associated with non-homogenous flower-like motility pattern on soft agar. In this study, we investigated the mechanisms explaining the motility reduction in the presence of polymyxin B. To do so, we took advantage of the flower-like motility pattern and performed an experimental evolution in the presence of polymyxin B in two strains of *V. cholerae, i*.*e*. O1 El Tor strain A1552 and O139 strain MO10. After sequencing, we identified mutations in genes involved in global regulation (*ihfA*), in outer membrane lipid asymmetry maintenance (*vacJ* and *mlaF*) and in cell wall synthesis (*dacB*).

## Material and methods

### Bacterial strains and growth conditions

Two *V. cholerae* strains have been used in this study: A1552 (O1, El Tor, Inaba) and MO10 (O139), initially isolated from patients in Peru and India, respectively [17, 18]. Bacterial growth was performed in LB media supplemented with 25 µg/mL of polymyxin B (Polymyxin B sulfate, Millipore) when indicated, as previously described [16]. Briefly, bacteria were grown overnight at 37°C under shaking condition in LB media followed by a dilution in fresh LB media. These cultures were then incubated in similar conditions to an OD of 0.3 and used to perform the motility assays.

### Motility assays

Motility assays were performed as previously described [16]. Briefly, bacteria were spotted on LB plates containing 0.3% agar (soft agar) and incubated for 12 to 48 hours at 37°C. The motility was evaluated by measuring the diameter of bacterial growth. Polymyxin B was added to the plate at a concentration of 25 µg/mL, a subinhibitory concentration that does not cause pore formation in the membrane [16], when indicated. Selection of variants was performed on technical triplicates. Experiments using variants were performed in biological triplicates. Statistical analyses were performed using a one-way ANOVA test (*p*<0.05).

### Sequencing, genome assembly and analysis

The DNA extraction of the bacterial strains was performed using Promega Wizard™ Genomic DNA Purification Kit according to the manufacturer guidelines. The wild-type strains MO10 and A1552 were sequenced by PacBio Sequel and Illumina MiSeq at the Génome Québec Innovation Center (Centre Hospitalier Universitaire Sainte-Justine, Montreal, Canada). The PacBio sequencing reads were assembled with Flye 2.8.1 [19]. The sequences corresponding to chromosomes I and II were circularized with various tools from the EMBOSS version 6.6.0 suite [20]. Illumina reads were subsequently used to polish assemblies with BWA version 0.7.17-r1188 [21], SAMtools version 1.12 [22] and Pilon version 1.23 [23]. The sequences were deposited in the NCBI database (<u>CP072847.1; CP072848.1; CP072849.1</u> and <u>CP072850.1</u>). The genomes are available on the NCBI biosample database: accession number SAMN18636297.

The DNA of the variants sequenced by Illumina MiSeq at the Génome Québec Innovation Center (McGill University, Montreal, Canada). The reads were filtered using fastp version 0.20.1 [24]. The mutations were identified and validated in relation to the genomes of the parental strains, beforehand annotated with Prokka version 1.14.5 [25], with both Snippy version 4.6.0 [26] and Breseq version 0.35.2 [27, 28].

### Sequences homologies, alignments and features analysis

*Escherichia coli* K12 MG1655 and *Vibrio cholerae* IhfA protein sequence identity and similarity were calculated with protein-protein BLAST [29, 30]. Alignment of two or multiple sequences were respectively performed using semi-global alignment “glocal” [31] and MUSCLE [32] from Snapgene^®^. Figure 5 were edited from Easyfig graphical sequence alignment [33], Figure 4 and Figure 6Figure ***7*** were edited from Snapgene^®^ alignments. Signal peptide was identified with protcompB (V9.0) [34] and SignalP 5.0 [35]. Putative -35 and -10 promoter box were identified with BPROM service from Softberry [36]. *Vibrio cholerae* gene and protein sequences homology was compared to *Escherichia coli* K12 MG1655 (<u>NC_000913.3</u>).

**Figure 1.**
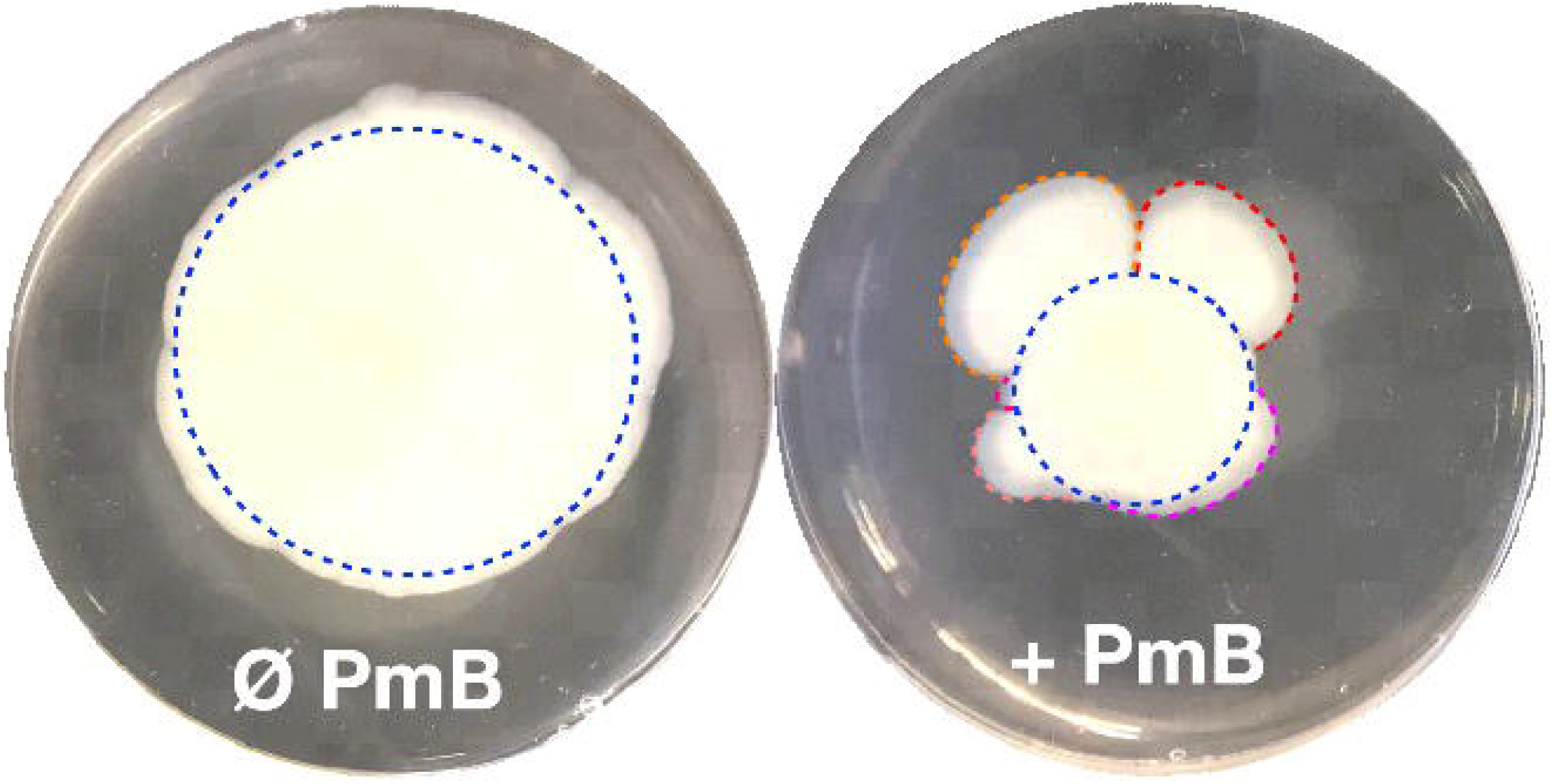
A flower-like pattern often appears in *Vibrio cholerae* motility test in presence of polymyxin B. Motility test on 0.3% soft agar in absence (Ø) or in presence (+) of 25 µg/mL polymyxin B (PmB). In the presence of PmB, motility pattern of *V. cholerae* are not symmetrical as we could expect (blue dotted circles) but show protrusions (red, orange, violet, salmon and pink curved lines). In absence of PmB, the motility pattern is much less heterogenous than in the presence of PmB. These “petals” suggest a spontaneous emergence of mutant more motile.

**Figure 2.**
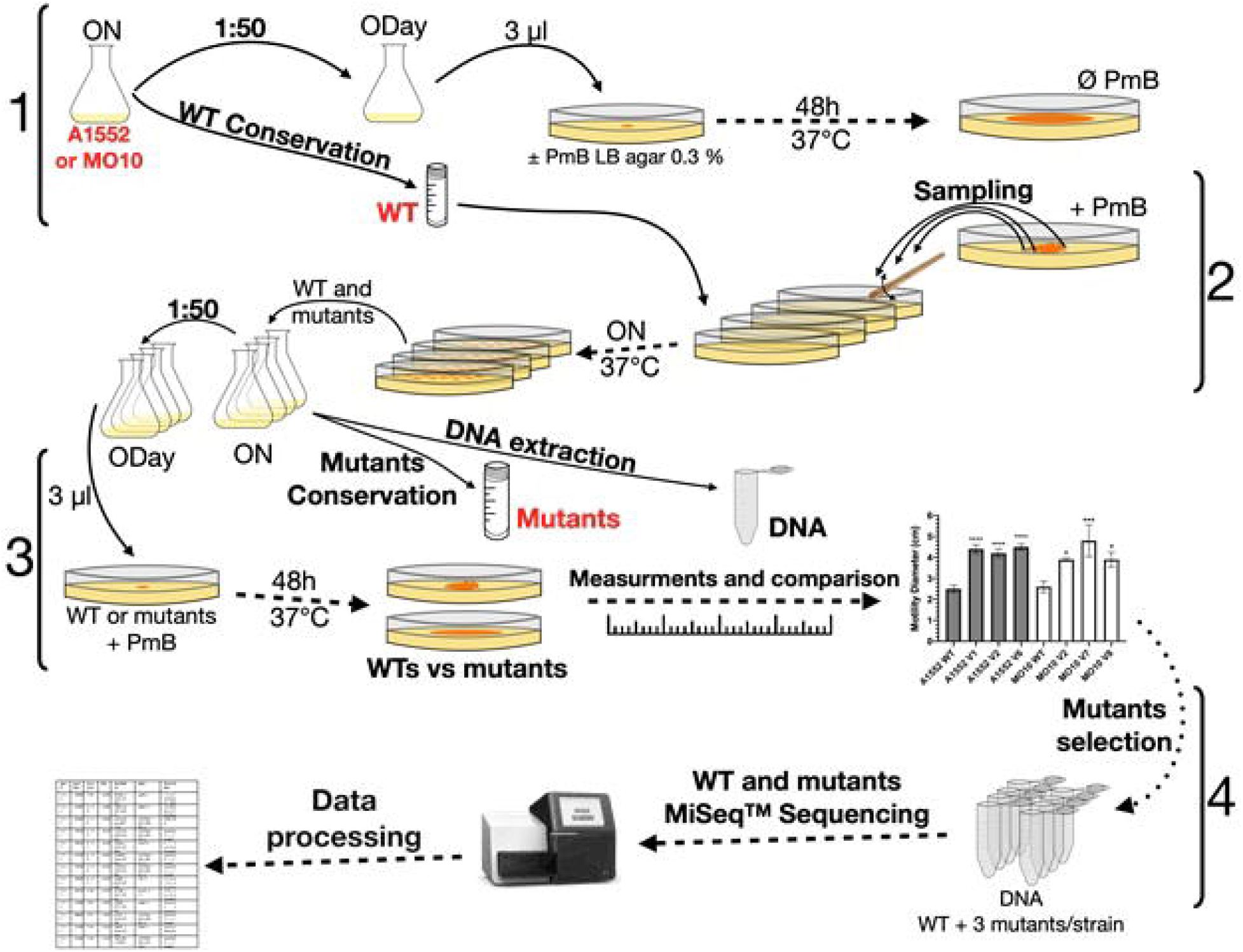
Directed evolution experimental design. In step 1, *V. cholerae* A1552 and MO10 strains were grown overnight, cells were conserved at this point as WT control and overday were started with 1:50 overnight culture. Then, 3 µl of both strains were spotted in technical triplicates on motility plates containing ± 25 µg/ml polymyxin B and incubated 48h at 37°C. In the second step, 4 samples were taken from the edge of different “petals” or protuberances of flower-like pattern of the different strains replicates then streaked on Petri dish. A colony from the different WT and putative mutant’s samples were grown overnight. Overnight cultures were diluted 1:50 to a fresh medium overday. A fraction of the mutants cultures were conserved at -80 °C. In the third step, the motility of the WT and selected variants were tested on plates containing PmB and grown at 37 °C for 48h. Motility diameters of the variants were measured and compared to their respective WT mother strains. In the last step, selected fast motility mutants and WT genomic DNA were sequenced using MiSeq™ technology. Data sequences were processed and analyzed. See material and methods for more information.

**Figure 3.**
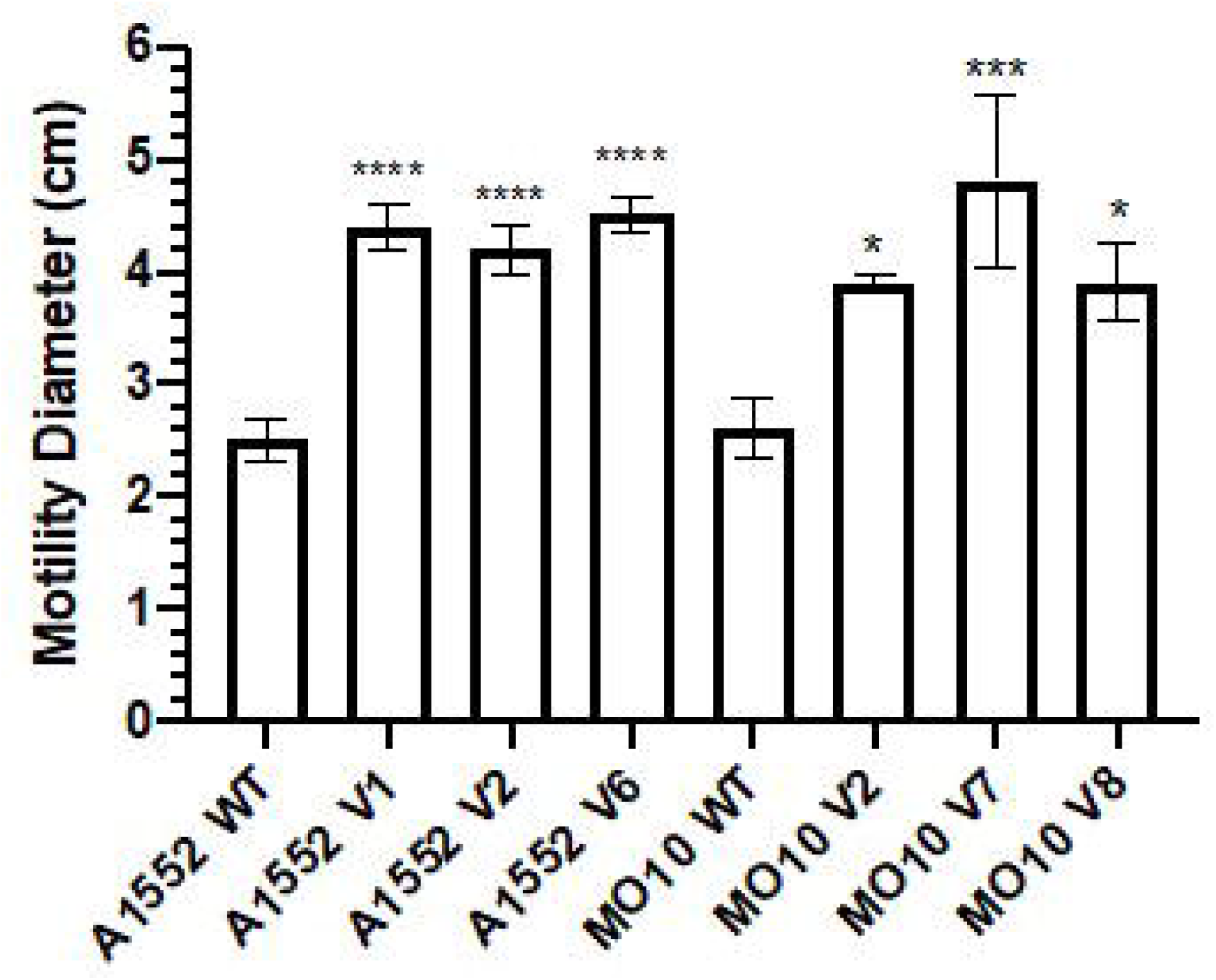
Selected sample from petals of A1552 and MO10 flower-like patterns present an increased motility. Swimming motility assay of A1552 and MO10 WT and corresponding variants. Exponential phase of growth aliquots of each strain was spotted in the center of a Petri dish containing LB agar 0.3% supplemented with 25 µg/mL of PmB. Motility diameters ± SD were measured after 48h incubation at 37 □C. Statistical significance between the variants motility diameter to their respective WT stains were tested by Dunnett one way ANOVA on biological triplicates for each strain. (* P<0.05) ; (** P<0.01) ; (*** P<0.001) ; (**** P<0,0001).

**Figure 4.**
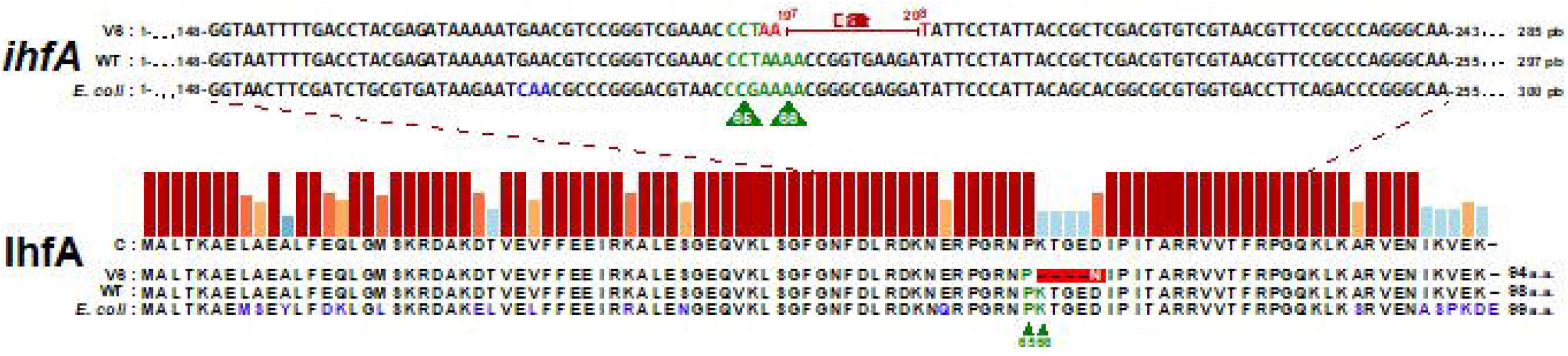
Loss of 12 nucleotides in *V. cholerae* A1552-V6 suggest a lost of an amino acid essential for IhfA function. Translated protein sequence alignment between *V. cholerae* A1552-V6 and WT strains and *E. coli* MG1655 strain. Sequence of *ihfA* gene [69] from *V. cholerae* A1552-V6 (V6) and *E. coli* MG1655 (*E. coli*) were translated and aligned to *V. cholerae* A1552 (WT) using MUSCLE [32] on Snapgene^®^. *V. cholerae* A1552 WT strain and *E. coli* IhfA proteins sequence shares 81% identity. The 2 amino acids P65 and K66 amino acids, essentials for IhfA function in E. coli [70], are also present in *Vibrio cholerae* A1552 WT strain. Modification in *ihfA* gene sequence leading to amino acid modifications in IhfA protein sequences are shown in blue. A1552-V6 variant had lost 12 nucleotides (197 to 208) in *ihfA* which induce 66 lysin replacement by asparagine (white letter) and loss of 4 amino acids.

**Figure 5.**
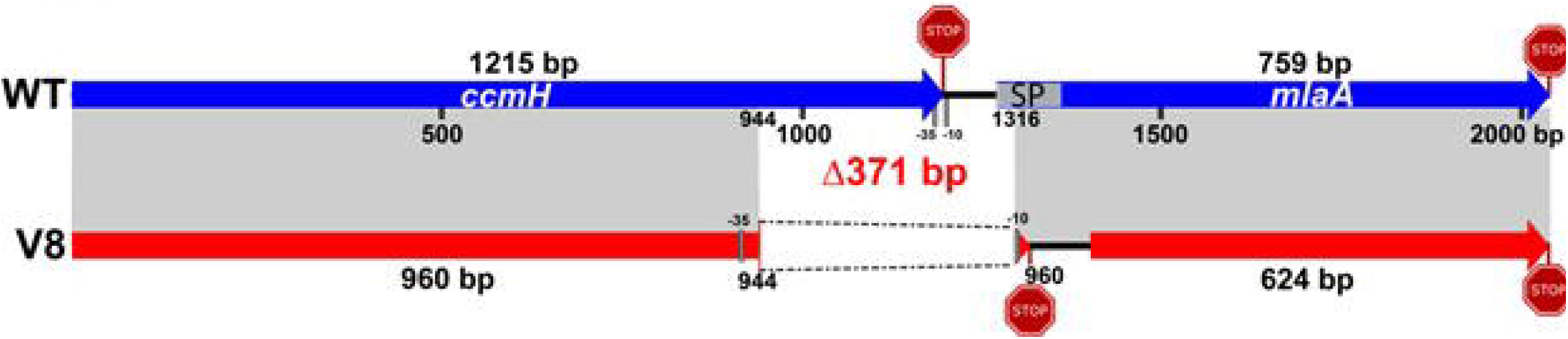
MO10-V8 has a deletion of 371 bp between *ccmH-1* and *vacJ (mlaA)* locus that probably cause a loss of function of CcmH and an absence of translation of *vacJ*. Schematic representation of *ccmH-1* (VchoM_01282) and *vacJ* (*mlaA*, VchoM_01281) locus alignment (in gray) between *V. cholerae* MO10 WT (blue) and MO10 V8 (red). The 371 bp deletion in V8 leads to the loss of ∼20% of *ccmH* 3’-end, including *ccmH* stop codon, and a frameshift leading to the emergence of a new stop codon in the *vacJ’s* 5’-ORF region. The *vacJ* gene has also lost important feature, like its -35 and -10 promoter regions and the first 30 coding nucleotides in outer membrane signal peptide (SP) sequence (nucleotide 1-51, codon 1-17). In V8 mutant, MlaA first AUG codon is compromised, the new predicted ORF starts 123 nucleotides (41 codons) forward. We detected new putative -35 and -10 promoter regions for *vacJ* in V8 but the signal peptide has been deleted. Figure were edited from Easyfig graphical sequence alignment [33]. Signal peptide was confirmed with protcompB (V9.0) [34] and SignalP 5.0 [35]. Putative -35 and -10 box for *vacJ \** at 923 and 944 using BPROM service from Softberry [36], see Figure S1.

**Figure 6.**
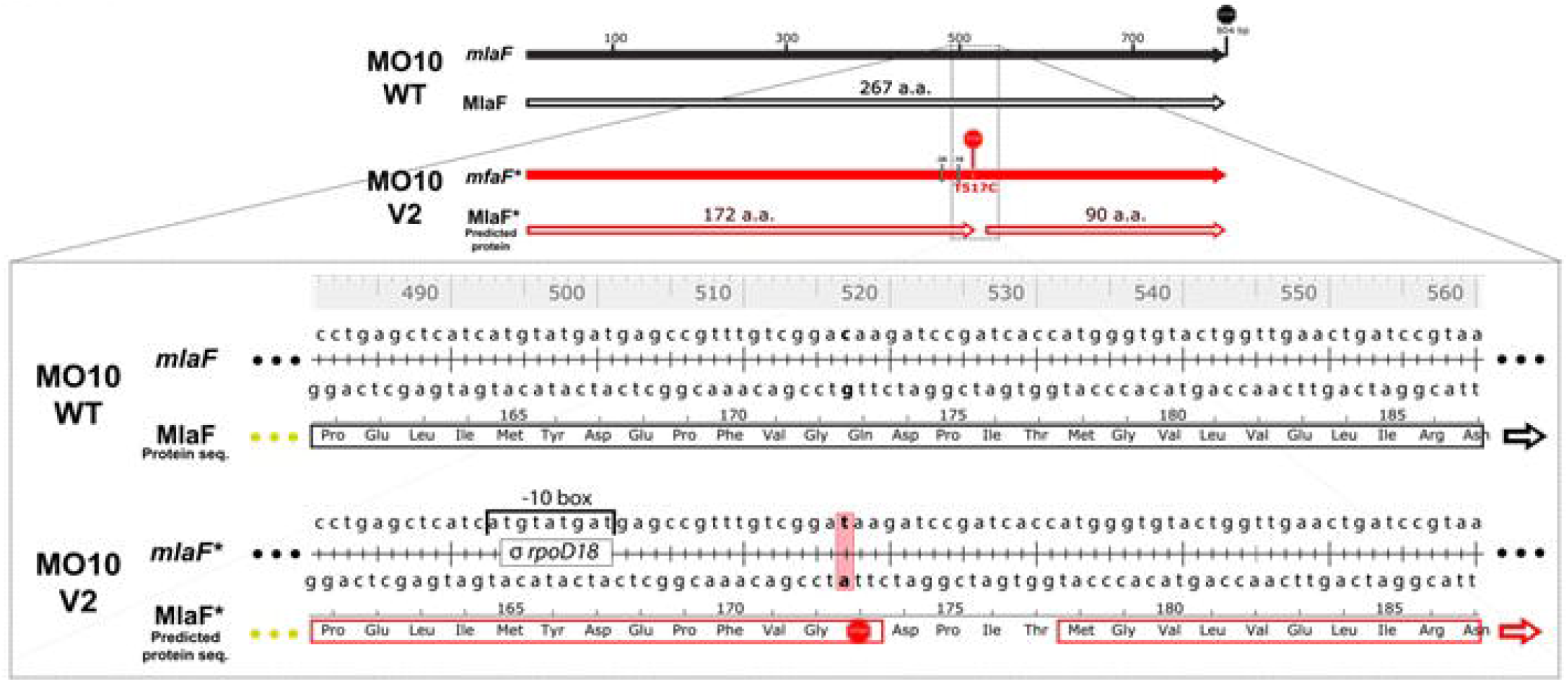
Non-sens mutation in *mlaF* results in the loss of 26% of original product. MiSeq sequencing results alignment of MO10 WT *mlaF* gene and MlaF protein sequence and MO10-V2 (V2) corresponding mutated gene (*mlaF*\*) and predicted protein sequence (MlaF*). Transversion of 517 thymine to cytosine in MO10-V2 resulted in a premature stop codon (codon 173). Compared to the WT MO10-V2 MlaF* has lost 95 amino acids (a. a.) corresponding to the ∼36% C-terminal end. A new ORF of 90 codons is created in the same reading frame, and a new promoter is predicted. Alignment were performed using MUSCLE [32] on Snapgene^®^ and promoter and were predicted using BPROM software [36], see Figure S2.

## Results

### Experimental evolution procedure

In some of our previous study, we observed a heterogenous motility pattern on soft agar in the presence of polymyxin B. Indeed, protuberances of various sizes were observed around the inner circle of bacteria (**Figure 1**). These protuberances were never observed in the control without polymyxin B (**Figure 1**). We hypothesize that these protuberances correspond to mutation in the bacterial chromosome which compensate the loss of motility in the presence of polymyxin B. Therefore, we performed an experimental evolution to select for adaptative variants that gain motility in presence of polymyxin B (**Figure 2**). To do so, we plated the wild-type strains on soft agar supplemented with 25 µg/mL of polymyxin B. After 48 hours, we sampled bacteria at the edge of the protuberances and streaked them on LB media to obtain isolated colonies. Two wild-type mother strains were used: *V. cholerae* O1 El Tor A1552 and O139 MO10. Eight sampling have been performed for each strain have been tested and one colony representative of each sample has been selected and conserved for a total of 16 variants: A1552 V1-8 and MO10-V1-8. Then, we tested the motility phenotype of this variants in presence of polymyxin B on soft agar. After 48h, 6 were selected based on their increased motility in presence of polymyxin B in comparison with the wild-type strains (**Figure 3**). From the 6 variants, 3 originated from *V. cholerae* A1552 (A1552-V1, A1552-V2 and A1552-V6) and 3 from MO10 (MO10-V1, MO10-V7 and MO10-V8). The motility of these variants are comparable to the motility of the wild-type strains in absence of polymyxin B [16] with a motility diameter of ∼4.3 cm on average (**Figure 3**). As expected, the wild-type strains in presence of polymyxin B demonstrated a reduced motility with a diameter ∼2.5 cm. Finally, the motility pattern was comparable with the control without polymyxin B displaying a more circular pattern (data not shown). Altogether, these results suggest that the variant have compensated the loss of motility observed in presence of polymyxin B.

### Identification of mutations

To determine the genes involved in the motility restoration of the variants in presence of polymyxin B, we performed a comparative genomic analysis. To do so, we first sequenced the genome of the wild-type strains using a long-read sequencing technology. The variants were sequenced using a MiSeq technology and compared it to the genome of the wild-type strains. The complete results of the comparative genomic analyses are presented in tables **Table S1** to **Table S5**.

The most significant mutation detected in A1552 variant occurs in the A1552-V6 variant and consist on a deletion of 12 nucleotides in *ihfA* (<u>VC1222</u>) gene (**Figure 4**). This mutation result in the loss of 4 amino-acids (66-69) and a mutation of one amino-acid in position 70 (D70N). A comparison of *ihfA* sequence in A1552 with its homologue in *E. coli* demonstrate that the sequence is highly conserved (94% similarity and 81% identity). Interestingly, the amino-acids responsible for IHF function in *E. coli* have been identified (P65 and K66) and are conserved in the wild-type sequence of A1552. Conversely, K66 is deleted in A1552-V6 (**Figure 4**). Therefore, it is possible that the protein remains transcribed but with an attenuated functionality.

Regarding MO10, two variants presented mutations in the Mla pathway, involved in phospholipid asymmetry maintenance. In MO10-V8, we noticed a deletion of 371 nucleotides in a region covering *ccmH* (<u>VC2050</u>) and *vacJ* (*mlaA*, <u>VC2048</u>) genes (**Figure 5**), while we observed a point mutation resulting in the insertion of a stop codon in *mlaF* (VC2520) sequence in MO10-V2 (**Figure 6**). Regarding *vacJ*, the 17 first codons were deleted, which result in the loss of the putative signal peptide for the secretion through the outer membrane as determined using SignalP 5.0 software [37]. Therefore, it is expected that both mutations in *vacJ* and *mlaF* induces a loss of function of these genes and proteins, and consequently of the Mla pathway. In *ccmH*, which encodes a c-type cytochrome [38, 39], the mutation result in the modification of 4 amino-acids and the deletion of the 85 subsequent amino-acids in C-terminal, which represent more than 20% of the sequence and the periplasmic domain of the protein.

In addition, we observed a 11 nucleotides deletion in *dacB* (<u>Vch1786_I0040</u>) in MO10-V2 in position 1013-1023 (**Figure 7**). This deletion result in a shift in the open reading frame and the emergence of a stop codon. In addition, DacB protein has lost ∼30% of its C-terminal residues. Therefore, it is very likely that *dacB* mutation in MO10-V2 result in a loss of function.

**Figure 7:**
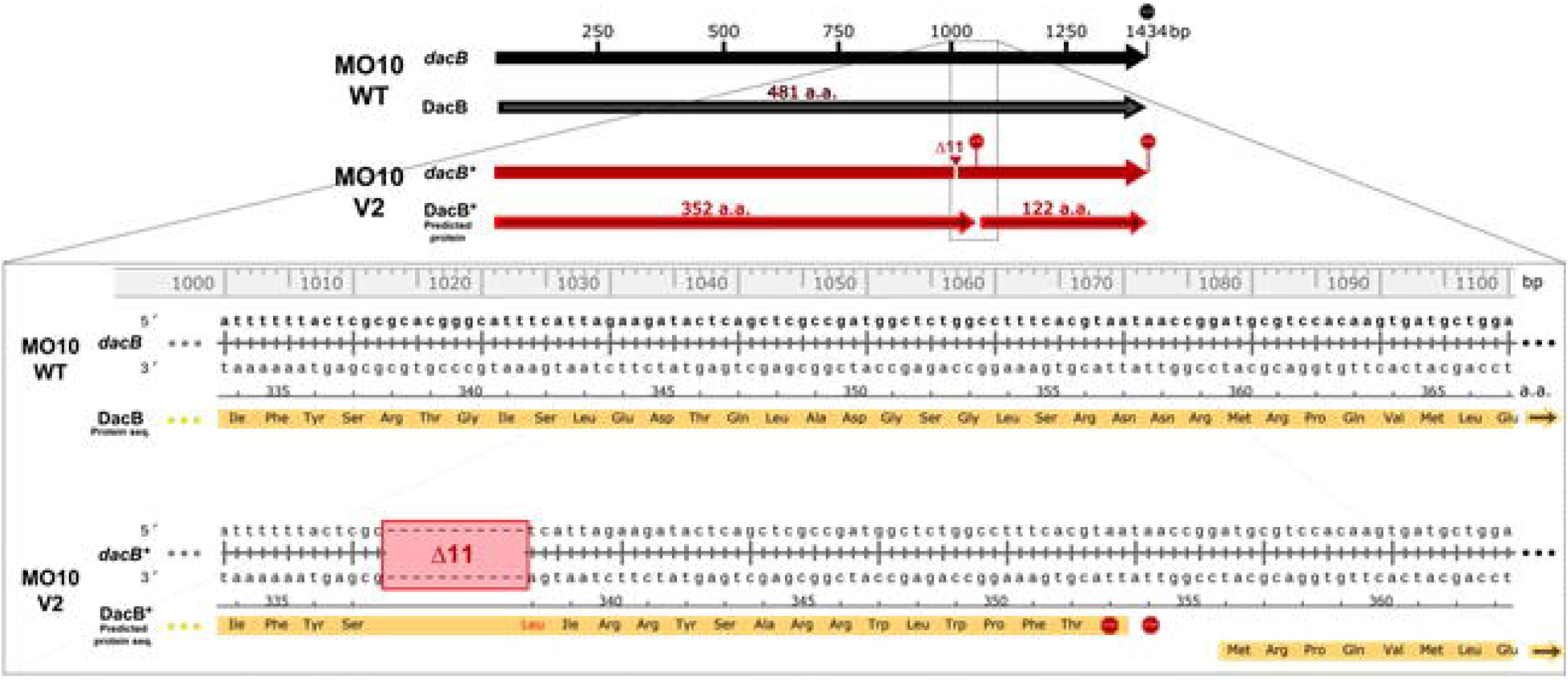
Deletion of 11 nucleotides in *dacB* in MO10-V2. MiSeq sequencing results alignment of MO10 WT *dacB* gene and DacB protein sequenc and MO10-V2 (V2) corresponding mutated gene (*dacB*\*) and predicted protein sequence (DacB*). MO10-V2 had lost 11 nucleotides (1013 to1023) resulting in a frame shift, causing modification of 16 amino acids, emergence of two early stop codons (STOP signs) and a new ORF starting at codon 360 of WT sequence. Alignment were performed using MUSCLE [32] on Snapgene^®^. Promoter were predicted using BPROM software [36], see Figure S3.

## Discussion

To identify genes important for motility in the presence of sub-inhibitory concentration of polymyxin B, we performed a short experimental evolution protocol. Experimental evolution has been widely used to validate evolutionary theories and to identify genes involved in responses to various stresses in bacteria [40]. In the latter case, *de novo* mutations driven by stresses led to the identification of genes important for survival in presence of oxidative stress and antibiotics, or during a temperature shift [41, 42]. In this study, we identified mutations in genes potentially important for motility in the presence of polymyxin B in two different strains of *V. cholerae*: A1552 (O1 El Tor) and MO10 (O139): *vacJ*-*ccmH mlaF, dacB* and *ihfA*. These mutations have been identified in three variants, two in MO10 and one in A1552.

In our previous study, we demonstrated that the loss of motility in the presence of polymyxin B was due to a high proportion of aflagellated bacteria [16]. The absence of flagella was not resulting from a repression of the flagellins genes, but most likely to either a default in the flagellum assembly or a flagella miss-anchoring. In any cases, the absence of flagella is related to envelope perturbations due to polymyxin B, while no pores in the membrane were observed [16]. Therefore, it was not surprising to identify in our analysis genes involved in the maintenance of the cell envelope integrity. It is the case of *vacJ* (*mlaA*) and *mlaF*, which belong to the maintenance of lipid assymetry (Mla) pathway. VacJ is homologue to MlaA, an alpha-helical outer membrane lipoprotein, while MlaF is an ATPAse associated to the inner membrane in complex with MlaD and MlaE. MlaC and MlaB are periplasmic and cytoplasmic proteins, respectively [43]. It has been previously described in *E. coli* that the Mla pathway is essential for preserving the outer membrane lipid asymmetry by preventing phospholipid accumulation in the outer leaflet through an anterograde phospholipid transport [43, 44]. The Mla pathway is important for antimicrobial tolerance in several Gram-negative bacteria, including *Burkholderia* and *Pseudomonas* [45, 46]. In *Vibrio cholerae* and other Gram-negative pathogens, it has been demonstrated that the Mla pathway is involved in membrane vesicles biogenesis and is important for serum resistance [47]. The authors proposed that the hypervesiculation mediated by the Mla pathway is a conserved bacterial mechanism to prevent phospholipid accumulation in the outer membrane [47]. It has been demonstrated that polymyxin B disturb the outer membrane and stiffen the phospholipid bilayer [48]. Thus, it is reasonable to hypothesize that the loss of *vacJ* or *mlaF* in presence of subinhibitory concentrations of polymyxin B might have a significant effect on outer membrane fluidity through vesiculation. Since the increased ratio of aflagellated bacteria is likely due to membrane perturbation [16], it is possible that the mutation in the Mla pathway observed in this study participate in stabilizing the membrane integrity, which would result in the preservation of the flagella and restored motility.

The deletion identified in MO10-V8 in *vacJ* also affect the sequence of the adjacent gene *ccmH*, which encodes for a c-type cytochrome. Cytochromes are important in bacteria metabolism as they function as electron transfer proteins. C-type cytochromes are characterized by the covalent liaison of the polypeptide with an heme [49]. In MO10-V8, the CXXCH motif, present in the periplasmic domain of the cytochromes c and essential for the heme binding [50] has been deleted has part of the 371 nucleotide deletion. Therefore, it is likely that CcmH has lost its function in this variant. In *V. cholerae*, there are 14 proteins containing a CXXCH motif and a signal peptide in N-terminal for the association with the membrane [39]. Among them, 7 have been characterized for their function in respiration in aerobic and anaerobic conditions and one is a peroxidase probably involved in oxidative stress response [39]. CcmH is part of the 6 other c-types cytochromes that are clustered together (*ccmA-I*) on *V. cholerae* chromosome and are not required for growth under aerobic conditions in rich media [39]. To our knowledge, there is no evidence that *ccmH* is important for motility or membrane integrity, and its role in bacterial metabolism is probably very limited in our experimental conditions. Therefore, the implication of the deletion that covers both *ccmH* and *vacJ* sequences in bacterial motility recovery in the presence of polymyxin B most probably rely on the inactivation of *vacJ*.

In another variant of the O139 strain MO10 strain, a mutation has been observed in *dacB*, which encodes PBP4, a low molecular weight cell wall-synthesizing enzymes/penicillin-binding proteins (PBPs) involved in peptidoglycan biosynthesis pathway [51]. In *Vibrio parahaemolyticus, dacB* has a role in the formation of abnormal cell shape during the transition to a viable but not culturable state [52]. In *E. coli*, low molecular weight PBPs also have a role in cell shape maintenance [53]. The *dacB* mutant cells are wider and display localized transparent bulges at the poles [54]. The authors also demonstrated that an accumulation of soluble peptidoglycan occurs in a *dacB* mutant. Additionally, it has been demonstrated that peptidoglycan remodelling, especially the alteration of the peptidoglycan-outer membrane cross linking, regulates the formation of membrane vesicles [55]. This alteration involves the endopeptidase activities of PBP4 and another protein, Spr, under the regulation by NlpI. Altogether, the roles of *dacB* in cell shape maintenance, peptidoglycan biosynthesis and vesicles formation, pinpoint its importance in cell wall remodelling that might be important for the stability of the flagella in *V. cholerae*.

The only mutation identified in the O1 El Tor strain A1552 is a deletion of 12 nucleotides in *ihfA. ihfA* encodes IfhA, one the Integration Host Factors (IHF) subunit together with IhfB. IHF is a histone-like DNA-binding protein that regulate many functions in bacterial cells, including transcription, replication and virulence [56]. In *V. cholerae*, IHF is essential for conjugation and virulence gene expression [57]. The deletion of 12 nucleotides likely result in an attenuation of IHF function since one of the two essential amino acids for DNA binding has been deleted. It has been demonstrated in a *V. cholerae* O1 El tor that IHF bind to *rpoN* and *flrA*, two major regulator of flagella genes expression [58]. The binding of IHF to *rpoN* and *flrA* promotors regions leads to a limitation of H-NS binding to these regions [58]. The role of H-NS as a negative regulator of *rpoN* and *flrA* has previously been described [59]. Thus, an IHF inactivation would lead to the binding of H-NS and eventually to the repression of the flagella related genes. However, we previously demonstrated that the expression of the flagellin subunits is not reduced in presence of polymyxin B [16]. Therefore, it is possible that the IHF regulation of the flagellin in the presence of polymyxin B is minimal. Since IHF binding sites have been identified all over the bacterial genomes [60, 61], it is reasonable to hypothesize that the mechanism behind the gain in motility in presence of polymyxin B involving *ihfA* mutation might not be related to the flagella itself. As stated above, the loss of motility in presence of polymyxin B is likely due to a structural defect in the flagella or in the cell envelop. It has been recently demonstrated that a deletion of *ihfA* in *Dickeya*, a bacteria belonging to the *Enterobactariaceae*, result in modifications of the cell envelope [60]. In addition, the expression of major porins of *E. coli* are controlled by IHF [62, 63]. In *Salmonella enterica*, an *ihfA* mutation induces the expression of genes involved in peptidoglycan and lipopolysaccharides biosynthesis [64]. Therefore, similarly to the Mla pathway and *dacB*, a mutation in *ihfA* might influence the expression of genes that strengthen the bacterial envelop, leading to the retention of the flagella in presence of polymyxin B.

Regarding the other variants that have been sequenced, no significant genetic mutations were observed. It is thus possible that the gain of motility observed in these variants is due to a transcriptional or an epigenetic regulation of gene expression. An increasing number of evidences suggest that epigenetic regulation through methylation is an important process for gene expression regulation, including antibiotic resistance and virulence [65, 66]. Regarding flagella, a methylation site has been reported in the *flh* operon in *E. coli* [67]. In addition, it has been demonstrated that the expression of genes involved in membrane integrity are also highly regulated, including through small regulatory RNAs [68]. Therefore, it is possible that the increase motility observed in these variants is due to a modulation in the expression of genes involved in flagella structure or in membrane integrity maintenance.

## References

1. Harris JB, LaRocque RC, Qadri F, Ryan ET, Calderwood SB. Cholera. The Lancet. 2012;379(9835):2466–76. doi: 10.1016/s0140-6736(12)60436-x.

2. Faruque SM, Sack DA. Sack RB, Colwell RR, Takeda Y, Nair GB. Emergence and evolution of Vibrio cholerae O139. Proc Natl Acad Sci U S A. 2003;100(3):1304-9. Epub 2003/01/23. doi: 10.1073/pnas.0337468100. PubMed PMID: 12538850; PubMed Central PMCID: PMCPMC298768.

3. Morris JG. Infections due to non-O1/O139 Vibrio cholerae. 2011.

4. Josenhans C, Suerbaum S. The role of motility as a virulence factor in bacteria. Int J Med Microbiol. 2002;291(8):605-14. Epub 2002/05/15. doi: 10.1078/1438-4221-00173. PubMed PMID: 12008914.

5. Kearns DB. A field guide to bacterial swarming motility. Nat Rev Microbiol. 2010;8(9):634-44. Epub 2010/08/10. doi: 10.1038/nrmicro2405. PubMed PMID: 20694026; PubMed Central PMCID: PMCPMC3135019.

6. Echazarreta MA, Klose KE. Vibrio Flagellar Synthesis. Front Cell Infect Microbiol. 2019;9:131. Epub 2019/05/24. doi: 10.3389/fcimb.2019.00131. PubMed PMID: 31119103; PubMed Central PMCID: PMCPMC6504787.

7. Nakamura S, Minamino T. Flagella-Driven Motility of Bacteria. Biomolecules. 2019;9(7). Epub 2019/07/25. doi: 10.3390/biom9070279. PubMed PMID: 31337100; PubMed Central PMCID: PMCPMC6680979.

8. Kojima S, Yamamoto K, Kawagishi I, Homma M. The polar flagellar motor of Vibrio cholerae is driven by an Na+ motive force. J Bacteriol. 1999;181(6):1927-30. Epub 1999/03/12. doi: 10.1128/JB.181.6.1927-1930.1999. PubMed PMID: 10074090; PubMed Central PMCID: PMCPMC93596.

9. Magariyama Y, Sugiyama S, Muramoto K, Maekawa Y, Kawagishi I, Imae Y, et al. Very fast flagellar rotation. Nature. 1994;371(6500):752. Epub 1994/10/27. doi: 10.1038/371752b0. PubMed PMID: 7935835.

10. Klose KE, Mekalanos JJ. Differential regulation of multiple flagellins in Vibrio cholerae. J Bacteriol. 1998;180(2):303-16. Epub 1998/01/24. doi: 10.1128/JB.180.2.303-316.1998. PubMed PMID: 9440520; PubMed Central PMCID: PMCPMC106886.

11. Fuerst JA, Perry JW. Demonstration of lipopolysaccharide on sheathed flagella of Vibrio cholerae O:1 by protein A-gold immunoelectron microscopy. J Bacteriol. 1988;170(4):1488-94. Epub 1988/04/01. doi: 10.1128/jb.170.4.1488-1494.1988. PubMed PMID: 2450866; PubMed Central PMCID: PMCPMC210992.

12. Ghosh A, Paul K, Chowdhury R. Role of the histone-like nucleoid structuring protein in colonization, motility, and bile-dependent repression of virulence gene expression in Vibrio cholerae. Infect Immun. 2006;74(5):3060-4. Epub 2006/04/20. doi: 10.1128/IAI.74.5.3060-3064.2006. PubMed PMID: 16622251; PubMed Central PMCID: PMCPMC1459692.

13. Correa NE, Peng F, Klose KE. Roles of the regulatory proteins FlhF and FlhG in the Vibrio cholerae flagellar transcription hierarchy. J Bacteriol. 2005;187(18):6324-32. Epub 2005/09/15. doi: 10.1128/JB.187.18.6324-6332.2005. PubMed PMID: 16159765; PubMed Central PMCID: PMCPMC1236648.

14. Kusumoto A, Shinohara A, Terashima H, Kojima S, Yakushi T, Homma M. Collaboration of FlhF and FlhG to regulate polar-flagella number and localization in Vibrio alginolyticus. Microbiology (Reading). 2008;154(Pt 5):1390-9. Epub 2008/05/03. doi: 10.1099/mic.0.2007/012641-0. PubMed PMID: 18451048.

15. Correa NE, Barker JR, Klose KE. The Vibrio cholerae FlgM homologue is an anti-sigma28 factor that is secreted through the sheathed polar flagellum. J Bacteriol. 2004;186(14):4613-9. Epub 2004/07/03. doi: 10.1128/JB.186.14.4613-4619.2004. PubMed PMID: 15231794; PubMed Central PMCID: PMCPMC438600.

16. Giacomucci S, Cros CD, Perron X, Mathieu-Denoncourt A, Duperthuy M. Flagella-dependent inhibition of biofilm formation by sub-inhibitory concentration of polymyxin B in Vibrio cholerae. PLoS One. 2019;14(8):e0221431. Epub 2019/08/21. doi: 10.1371/journal.pone.0221431. PubMed PMID: 31430343; PubMed Central PMCID: PMC6701800.

17. Allue-Guardia A, Echazarreta M, Koenig SSK, Klose KE, Eppinger M. Closed Genome Sequence of Vibrio cholerae O1 El Tor Inaba Strain A1552. Genome Announc. 2018;6(9). Epub 2018/03/03. doi: 10.1128/genomeA.00098-18. PubMed PMID: 29496831; PubMed Central PMCID: PMCPMC5834340.

18. O’Shea YA, Reen FJ, Quirke AM, Boyd EF. Evolutionary genetic analysis of the emergence of epidemic Vibrio cholerae isolates on the basis of comparative nucleotide sequence analysis and multilocus virulence gene profiles. J Clin Microbiol. 2004;42(10):4657-71. Epub 2004/10/09. doi: 10.1128/JCM.42.10.4657-4671.2004. PubMed PMID: 15472325; PubMed Central PMCID: PMCPMC522369.

19. Kolmogorov M, Yuan J, Lin Y, Pevzner PA. Assembly of long, error-prone reads using repeat graphs. Nat Biotechnol. 2019;37(5):540-6. Epub 2019/04/03. doi: 10.1038/s41587-019-0072-8. PubMed PMID: 30936562.

20. Rice P, Longden I, Bleasby A. EMBOSS: The European Molecular Biology Open Software Suite. Trends in Genetics. 2000;16(6):276–7. doi: 10.1016/s0168-9525(00)02024-2.

21. Li H. Aligning sequence reads, clone sequences and assembly contigs with BWA-MEM. arXiv. 2013.

22. Li H, Handsaker B, Wysoker A, Fennell T, Ruan J, Homer N, et al. The Sequence Alignment/Map format and SAMtools. Bioinformatics. 2009;25(16):2078-9. Epub 2009/06/10. doi: 10.1093/bioinformatics/btp352. PubMed PMID: 19505943; PubMed Central PMCID: PMCPMC2723002.

23. Walker BJ, Abeel T, Shea T, Priest M, Abouelliel A, Sakthikumar S, et al. Pilon: an integrated tool for comprehensive microbial variant detection and genome assembly improvement. LoS One. 2014;9(11):e112963. Epub 2014/11/20. doi: 10.1371/journal.pone.0112963. PubMed PMID: 25409509; PubMed Central PMCID: PMCPMC4237348.

24. Chen S, Zhou Y, Chen Y, Gu J. fastp: an ultra-fast all-in-one FASTQ preprocessor. Bioinformatics. 2018;34(17):i884-i90. Epub 2018/11/14. doi: 10.1093/bioinformatics/bty560. PubMed PMID: 30423086; PubMed Central PMCID: PMCPMC6129281.

25. Seemann T. Prokka: rapid prokaryotic genome annotation. Bioinformatics. 2014;30(14):2068-9. Epub 2014/03/20. doi: 10.1093/bioinformatics/btu153. PubMed PMID: 24642063.

26. Snippy: fast bacterial variant calling from NGS reads. In: Seemann T, editor. 4.0 ed2015.

27. Barrick JE, Lenski RE. Genome-wide mutational diversity in an evolving population of Escherichia coli. Cold Spring Harb Symp Quant Biol. 2009;74:119-29. Epub 2009/09/25. doi: 10.1101/sqb.2009.74.018. PubMed PMID: 19776167; PubMed Central PMCID: PMCPMC2890043.

28. Barrick JE, Yu DS, Yoon SH, Jeong H, Oh TK, Schneider D, et al. Genome evolution and adaptation in a long-term experiment with Escherichia coli. Nature. 2009;461(7268):1243-7. Epub 2009/10/20. doi: 10.1038/nature08480. PubMed PMID: 19838166.

29. Altschul SF, Madden TL, Schaffer AA, Zhang J, Zhang Z, Miller W, et al. Gapped BLAST and PSI-BLAST: a new generation of protein database search programs. Nucleic Acids Res. 1997;25(17):3389-402. Epub 1997/09/01. doi: 10.1093/nar/25.17.3389. PubMed PMID: 9254694; PubMed Central PMCID: PMCPMC146917.

30. Altschul SF, Wootton JC, Gertz EM, Agarwala R, Morgulis A, Schaffer AA, et al. Protein database searches using compositionally adjusted substitution matrices. FEBS J. 2005;272(20):5101-9. Epub 2005/10/13. doi: 10.1111/j.1742-4658.2005.04945.x. PubMed PMID: 16218944; PubMed Central PMCID: PMCPMC1343503.

31. Brudno M, Malde S, Poliakov A, Do CB, Couronne O, Dubchak I, et al. Glocal alignment: finding rearrangements during alignment. Bioinformatics. 2003;19 Suppl 1:i54-62. Epub 2003/07/12. doi: 10.1093/bioinformatics/btg1005. PubMed PMID: 12855437.

32. Edgar RC. MUSCLE: multiple sequence alignment with high accuracy and high throughput. Nucleic Acids Res. 2004;32(5):1792-7. Epub 2004/03/23. doi: 10.1093/nar/gkh340. PubMed PMID: 15034147; PubMed Central PMCID: PMCPMC390337.

33. Sullivan MJ, Petty NK, Beatson SA. Easyfig: a genome comparison visualizer. Bioinformatics. 2011;27(7):1009-10. Epub 2011/02/01. doi: 10.1093/bioinformatics/btr039. PubMed PMID: 21278367; PubMed Central PMCID: PMCPMC3065679.

34. ProtCompB - Prediction sub-cellular protein localization [20 avril 2021]. Available from: http://www.softberry.com/berry.phtml?topic=pcompb&group=programs&subgroup=proloc.

35. Nielsen H, Tsirigos KD, Brunak S, von Heijne G. A Brief History of Protein Sorting Prediction. Protein J. 2019;38(3):200-16. Epub 2019/05/24. doi: 10.1007/s10930-019-09838-3. PubMed PMID: 31119599; PubMed Central PMCID: PMCPMC6589146.

36. V. Solovyev, Salamov A. Automatic Annotation of Microbial Genomes and Metagenomic Sequences 2011. In: Metagenomics and its applications in agriculture, biomedicine, and environmental studies [Internet]. Hauppauge, N.Y.: Nova Science Publisher’s; [61-78]. Available from: http://www.softberry.com/berry.phtml?topic=bprom&group=programs&subgroup=gfindb; https://public.ebookcentral.proquest.com/choice/publicfullrecord.aspx?p=3020875.

37. Almagro Armenteros JJ, Tsirigos KD, Sonderby CK, Petersen TN, Winther O, Brunak S, et al. SignalP 5.0 improves signal peptide predictions using deep neural networks. Nat Biotechnol. 2019;37(4):420-3. Epub 2019/02/20. doi: 10.1038/s41587-019-0036-z. PubMed PMID: 30778233.

38. Thony-Meyer L. Biogenesis of respiratory cytochromes in bacteria. Microbiol Mol Biol Rev. 1997;61(3):337-76. Epub 1997/09/18. doi: 1092-2172/97/$04.00?0. PubMed PMID: 9293186; PubMed Central PMCID: PMCPMC232615.

39. Braun M, Thony-Meyer L. Cytochrome c maturation and the physiological role of c-type cytochromes in Vibrio cholerae. J Bacteriol. 2005;187(17):5996-6004. Epub 2005/08/20. doi: 10.1128/JB.187.17.5996-6004.2005. PubMed PMID: 16109941; PubMed Central PMCID: PMCPMC1196146.

40. McDonald MJ. Microbial Experimental Evolution - a proving ground for evolutionary theory and a tool for discovery. EMBO Rep. 2019;20(8):e46992. Epub 2019/07/25. doi: 10.15252/embr.201846992. PubMed PMID: 31338963; PubMed Central PMCID: PMCPMC6680118.

41. Justin T, Connor A. O, Joon Ho P, Anand V. S, Patrick V. Phaneuf, Laurance Y, et al. Experimental evolution reveals the genetic basis and systems biology of superoxide stress tolerance. 2019. doi: 10.1101/749887.

42. Hoffmann AA, Hercus MJ. Environmental Stress as an Evolutionary Force. BioScience. 2000;50(3). doi: 10.1641/0006-3568(2000)050[0217:Esaaef]2.3.Co;2.

43. Du Toit A. Phospholipid export from the inside out. Nat Rev Microbiol. 2019;17(9):528. Epub 2019/07/11. doi: 10.1038/s41579-019-0239-9. PubMed PMID: 31289381.

44. Malinverni JC, Silhavy TJ. An ABC transport system that maintains lipid asymmetry in the gramnegative outer membrane. Proc Natl Acad Sci U S A. 2009;106(19):8009-14. Epub 2009/04/23. doi: 10.1073/pnas.0903229106. PubMed PMID: 19383799; PubMed Central PMCID: PMCPMC2683108.

45. Bernier SP, Son S, Surette MG. The Mla Pathway Plays an Essential Role in the Intrinsic Resistance of Burkholderia cepacia Complex Species to Antimicrobials and Host Innate Components. J Bacteriol. 2018;200(18). Epub 2018/07/11. doi: 10.1128/JB.00156-18. PubMed PMID: 29986943; PubMed Central PMCID: PMCPMC6112004.

46. Munguia J, LaRock DL, Tsunemoto H, Olson J, Cornax I, Pogliano J, et al. The Mla pathway is critical for Pseudomonas aeruginosa resistance to outer membrane permeabilization and host innate immune clearance. J Mol Med (Berl). 2017;95(10):1127-36. Epub 2017/08/28. doi: 10.1007/s00109-017-1579-4. PubMed PMID: 28844103; PubMed Central PMCID: PMCPMC5671890.

47. Roier S, Zingl FG, Cakar F, Durakovic S, Kohl P, Eichmann TO, et al. A novel mechanism for the biogenesis of outer membrane vesicles in Gram-negative bacteria. Nat Commun. 2016;7:10515. Epub 2016/01/26. doi: 10.1038/ncomms10515. PubMed PMID: 26806181; PubMed Central PMCID: PMCPMC4737802.

48. Fu L, Wan M, Zhang S, Gao L, Fang W. Polymyxin B Loosens Lipopolysaccharide Bilayer but Stiffens Phospholipid Bilayer. Biophys J. 2020;118(1):138-50. Epub 2019/12/10. doi: 10.1016/j.bpj.2019.11.008. PubMed PMID: 31812355; PubMed Central PMCID: PMCPMC6950770.

49. Kranz RG, Richard-Fogal C, Taylor JS, Frawley ER. Cytochrome c biogenesis: mechanisms for covalent modifications and trafficking of heme and for heme-iron redox control. Microbiol Mol Biol Rev. 2009;73(3):510-28, Table of Contents. Epub 2009/09/02. doi: 10.1128/MMBR.00001-09. PubMed PMID: 19721088; PubMed Central PMCID: PMCPMC2738134.

50. Barker PD, Ferguson SJ. Still a puzzle: why is haem covalently attached in c-type cytochromes? Structure. 1999;7(12):R281–R90. doi: 10.1016/s0969-2126(00)88334-3.

51. Typas A, Banzhaf M, Gross CA, Vollmer W. From the regulation of peptidoglycan synthesis to bacterial growth and morphology. Nat Rev Microbiol. 2011;10(2):123-36. Epub 2011/12/29. doi: 10.1038/nrmicro2677. PubMed PMID: 22203377; PubMed Central PMCID: PMCPMC5433867.

52. Hung WC, Jane WN, Wong HC. Association of a D-alanyl-D-alanine carboxypeptidase gene with the formation of aberrantly shaped cells during the induction of viable but nonculturable Vibrio parahaemolyticus. Appl Environ Microbiol. 2013;79(23):7305-12. Epub 2013/09/24. doi: 10.1128/AEM.01723-13. PubMed PMID: 24056454; PubMed Central PMCID: PMCPMC3837741.

53. Nelson DE, Young KD. Contributions of PBP 5 and DD-carboxypeptidase penicillin binding proteins to maintenance of cell shape in Escherichia coli. J Bacteriol. 2001;183(10):3055-64. Epub 2001/04/28. doi: 10.1128/JB.183.10.3055-3064.2001. PubMed PMID: 11325933; PubMed Central PMCID: PMCPMC95205.

54. Yang H, Lu X, Hu J, Chen Y, Shen W, Liu L. Boosting Secretion of Extracellular Protein by Escherichia coli via Cell Wall Perturbation. Appl Environ Microbiol. 2018;84(20). Epub 2018/08/12. doi: 10.1128/AEM.01382-18. PubMed PMID: 30097440; PubMed Central PMCID: PMCPMC6182897.

55. Schwechheimer C, Rodriguez DL, Kuehn MJ. NlpI-mediated modulation of outer membrane vesicle production through peptidoglycan dynamics in Escherichia coli. Microbiologyopen. 2015;4(3):375-89. Epub 2015/03/11. doi: 10.1002/mbo3.244. PubMed PMID: 25755088; PubMed Central PMCID: PMCPMC4475382.

56. Freundlich M, Ramani N, Mathew E, Sirko A, Tsui P. The role of integration host factor in gene expression in Escherichia coli. Mol Microbiol. 1992;6(18):2557-63. Epub 1992/09/01. doi: 10.1111/j.1365-2958.1992.tb01432.x. PubMed PMID: 1447969.

57. McLeod SM, Burrus V, Waldor MK. Requirement for Vibrio cholerae integration host factor in conjugative DNA transfer. J Bacteriol. 2006;188(16):5704-11. Epub 2006/08/04. doi: 10.1128/JB.00564-06. PubMed PMID: 16885438; PubMed Central PMCID: PMCPMC1540070.

58. Wang H, Ayala JC, Benitez JA, Silva AJ. Interaction of the histone-like nucleoid structuring protein and the general stress response regulator RpoS at Vibrio cholerae promoters that regulate motility and hemagglutinin/protease expression. J Bacteriol. 2012;194(5):1205-15. Epub 2011/12/24. doi: 10.1128/JB.05900-11. PubMed PMID: 22194453; PubMed Central PMCID: PMCPMC3294804.

59. Prouty MG, Correa NE, Klose KE. The novel sigma54-and sigma28-dependent flagellar gene transcription hierarchy of Vibrio cholerae. Mol Microbiol. 2001;39(6):1595-609. Epub 2001/03/22. doi: 10.1046/j.1365-2958.2001.02348.x. PubMed PMID: 11260476.

60. Reverchon S, Meyer S, Forquet R, Hommais F, Muskhelishvili G, Nasser W. The nucleoid-associated protein IHF acts as a ‘transcriptional domainin’ protein coordinating the bacterial virulence traits with global transcription. Nucleic Acids Res. 2021;49(2):776-90. Epub 2020/12/19. doi: 10.1093/nar/gkaa1227. PubMed PMID: 33337488; PubMed Central PMCID: PMCPMC7826290.

61. Prieto AI, Kahramanoglou C, Ali RM, Fraser GM, Seshasayee AS, Luscombe NM. Genomic analysis of DNA binding and gene regulation by homologous nucleoid-associated proteins IHF and HU in Escherichia coli K12. Nucleic Acids Res. 2012;40(8):3524-37. Epub 2011/12/20. doi: 10.1093/nar/gkr1236. PubMed PMID: 22180530; PubMed Central PMCID: PMCPMC3333857.

62. Ramani N, Huang L, Freundlich M. In vitro interactions of integration host factor with the ompF promoter-regulatory region of Escherichia coli. Mol Gen Genet. 1992;231(2):248-55. Epub 1992/01/01. doi: 10.1007/BF00279798. PubMed PMID: 1736095.

63. Huang L, Tsui P, Freundlich M. Integration host factor is a negative effector of in vivo and in vitro expression of ompC in Escherichia coli. J Bacteriol. 1990;172(9):5293-8. Epub 1990/09/01. doi: 10.1128/jb.172.9.5293-5298.1990. PubMed PMID: 2203749; PubMed Central PMCID: PMCPMC213192.

64. Mangan MW, Lucchini S, Danino V, Croinin TO, Hinton JC, Dorman CJ. The integration host factor (IHF) integrates stationary-phase and virulence gene expression in Salmonella enterica serovar Typhimurium. Mol Microbiol. 2006;59(6):1831-47. Epub 2006/03/24. doi: 10.1111/j.1365-2958.2006.05062.x. PubMed PMID: 16553887.

65. Ghosh D, Veeraraghavan B, Elangovan R, Vivekanandan P. Antibiotic Resistance and Epigenetics: More to It than Meets the Eye. Antimicrob Agents Chemother. 2020;64(2). Epub 2019/11/20. doi: 10.1128/AAC.02225-19. PubMed PMID: 31740560; PubMed Central PMCID: PMCPMC6985748.

66. Gaultney RA, Vincent AT, Lorioux C, Coppee JY, Sismeiro O, Varet H, et al. 4-Methylcytosine DNA modification is critical for global epigenetic regulation and virulence in the human pathogen Leptospira interrogans. Nucleic Acids Res. 2020;48(21):12102-15. Epub 2020/12/11. doi: 10.1093/nar/gkaa966. PubMed PMID: 33301041; PubMed Central PMCID: PMCPMC7708080.

67. Wang MX, Church GM. A whole genome approach to in vivo DNA-protein interactions in E. coli. Nature. 1992;360(6404):606-10. Epub 1992/12/10. doi: 10.1038/360606a0. PubMed PMID: 1334233.

68. Guillier M, Gottesman S. Remodelling of the Escherichia coli outer membrane by two small regulatory RNAs. Mol Microbiol. 2006;59(1):231-47. Epub 2005/12/20. doi: 10.1111/j.1365-2958.2005.04929.x. PubMed PMID: 16359331.

69. Kakkanat A, Phan MD, Lo AW, Beatson SA, Schembri MA. Novel genes associated with enhanced motility of Escherichia coli ST131. PLoS One. 2017;12(5):e0176290. Epub 2017/05/11. doi: 10.1371/journal.pone.0176290. PubMed PMID: 28489862; PubMed Central PMCID: PMCPMC5425062.

70. Lee EC, Hales LM, Gumport RI, Gardner JF. The isolation and characterization of mutants of the integration host factor (IHF) of Escherichia coli with altered, expanded DNA-binding specificities. The EMBO Journal. 1992;11(1):305–13. doi: 10.1002/j.1460-2075.1992.tb05053.x.

