## Supplementary file for "Experimental evolution of *Vibrio cholerae*: Identification of genes involved in motility in presence of polymyxin B"

**Table S1 Mutation accumulated in A1552 V1**

| **Chromosome** | **Position** | **Mutation** | **Frequency** | **Annotation** | **Gene name or locus ^a^** | **Description** |
| --- | --- | --- | --- | --- | --- | --- |
| Chr 2 | 528239 | T→G | 22,0% | K244Q (AAA→CAA) | *yejB ←* | Inner membrane ABC transporter permease protein YejB |
| Chr 1 | 559574 | A→G | 14,8% | G83G (GGA→GGG) | *dhbA →* | 2,3‑dihydro‑2,3‑dihydroxybenzoate dehydrogenase |
| Chr 1 | 85397 | G→A | 14,4% | noncoding (64/77 nt) | *J9265_00430 →* | tRNA‑Met |
| Chr 1 | 797412 | A→G | 13,7% | T1074A (ACG→GCG) | *J9265_03690 →* | hypothetical protein |
| Chr 1 | 2921010 | G→C | 13,3% | G292G (GGC→GGG) | *murP ←* | PTS system N‑acetylmuramic acid‑specific EIIBC component |
| Chr 1 | 797441 | A→G | 10,2% | E1083E (GAA→GAG) | *J9265_03690 →* | hypothetical protein |
| Chr 1 | 797417 | G→A | 9,7% | E1075E (GAG→GAA) | *J9265_03690 →* | hypothetical protein |
| Chr 1 | 585650 | A→G | 6,2% | E348G (GAA→GGA) | *citF →* | Citrate lyase alpha chain |
| Chr 1 | 2977196 | C→A | 5,7% | A710D (GCC→GAC) | *bepE_2 →* | Efflux pump membrane transporter BepE |
| Chr 1 | 1593654 | T→C | 5,3% | V289V (GTT→GTC) | *sapB →* | Putrescine export system permease protein SapB |
| Chr 2 | 888250 | C→T | 5,3% | A157V (GCG→GTG) | *J9265_18105 →* | hypothetical protein |
| Chr 1 | 2179501 | T→C | 5,2% | K302E (AAA→GAA) | *lpxB ←* | Lipid‑A‑disaccharide synthase |
| Chr 1 | 1263031 | C→A | 5,1% | S48I (AGC→ATC) | *ccoS ←* | cbb3-type cytochrome oxidase assembly protein CcoS |

a : Gene or locus name corresponding to *Vibrio cholerae* A1552 chromosome 1 [CP072847.1](https://www.ncbi.nlm.nih.gov/nuccore/CP072847.1) or chromosome 2 : [CP072848.1](https://www.ncbi.nlm.nih.gov/nuccore/CP072848.1).

**Table S2 Mutation accumulated in A1552 V2**

| **Chromosome** | **Position** | **Mutation** | **Frequency** | **Annotation** | **Gene name or locus ^a^** | **Description** |
| --- | --- | --- | --- | --- | --- | --- |
| Chr 1 | 85385 | C→T | 15,6% | noncoding (52/77 nt) | *J9265_00430 →* | tRNA‑Met |
| Chr 1 | 228686 | C→A | 12,6% | M292I (ATG→ATT) | *metK ←* | S‑adenosylmethionine synthase |
| Chr 1 | 310864 | C→T | 12,6% | intergenic (+13/‑44) | *J9265_01575 → / → J9265_01580* | tRNA‑Arg/tRNA‑Arg |
| Chr 1 | 1207946 | G→T | 9,1% | G164V (GGC→GTC) | *dnaK_2 →* | Chaperone protein DnaK |
| Chr 1 | 936910 | T→C | 8,5% | Y247Y (TAT→TAC) | *hisF →* | Imidazole glycerol phosphate synthase subunit HisF |
| Chr 1 | 1731673 | T→C | 7,6% | K109K (AAA→AAG) | *J9265_07800 ←* | hypothetical protein |
| Chr 1 | 2668056 | Δ1 bp | 6,6% | coding (642/2322 nt) | *yhgF →* | Protein YhgF |
| Chr 1 | 2729296 | A→G | 6,6% | M314T (ATG→ACG) | *ftsY_2 ←* | Signal recognition particle receptor FtsY |
| Chr 1 | 2602710 | G→T | 6,5% | Q2K (CAA→AAA) | *yibN ←* | putative protein YibN |
| Chr 1 | 2668058 | Δ1 bp | 6,5% | coding (644/2322 nt) | *yhgF →* | Protein YhgF |
| Chr 1 | 1349691 | G→T | 6,4% | A510D (GCT→GAT) | *J9265_06075 ←* | hypothetical protein |
| Chr 1 | 400889 | G→T | 6,2% | P50P (CCC→CCA) | *mepM_2 ←* | Murein DD‑endopeptidase MepM |
| Chr 1 | 2337536 | C→A | 6,0% | D26Y (GAC→TAC) | *mutT ←* | 8‑oxo‑dGTP diphosphatase |
| Chr 1 | 2473170 | C→A | 0,058 | E201* (GAG→TAG) | *rapA ←* | RNA polymerase‑associated protein RapA |
| Chr 1 | 95645 | G→C | 5,5% | G51A (GGT→GCT) | *hflC →* | Modulator of FtsH protease HflC |
| Chr 2 | 581386 | T→C | 5,3% | K534E (AAG→GAG) | *J9265_16790 ←* | hypothetical protein |
| Chr 2 | 664326 | G→T | 5,3% | L655F (TTG→TTT) | *yccS →* | Inner membrane protein YccS |
| Chr 1 | 486553 | G→A | 5,2% | R95C (CGC→TGC) | *bamD ←* | Outer membrane protein assembly factor BamD |
| Chr 1 | 2639649 | C→A | 5,1% | P86H (CCT→CAT) | *sodA →* | Superoxide dismutase [Mn] |

a : Gene or locus name corresponding to *Vibrio cholerae* A1552 chromosome 1 [CP072847.1](https://www.ncbi.nlm.nih.gov/nuccore/CP072847.1) or chromosome 2 : [CP072848.1](https://www.ncbi.nlm.nih.gov/nuccore/CP072848.1).

**Table S3 Mutation accumulated in A1552 V6**

| **Chromosome** | **Position** | **Mutation** | **Frequency** | **Annotation** | **Gene name or locus ^a^** | **Description** |
| --- | --- | --- | --- | --- | --- | --- |
| Chr 1 | 1029617 | Δ12 bp | 100,0% | coding (187‑198/297 nt) | *ihfA →* | Integration host factor subunit alpha |
| Chr 2 | 1048234 | G→T | 12,3% | A463E (GCA→GAA) | *J9265_18815 ←* | hypothetical protein |
| Chr 1 | 1458543 | T→A | 12,0% | D84E (GAT→GAA) | *macB_2 →* | Macrolide export ATP‑binding/permease protein MacB |
| Chr 1 | 2091693 | A→T | 6,3% | N74I (AAT→ATT) | *wrbA →* | NAD(P)H dehydrogenase (quinone) |
| Chr 1 | 1170344 | A→C | 6,0% | L1318F (TTA→TTC) | *rcsC_4 →* | Sensor histidine kinase RcsC |
| Chr 1 | 797108 | C→A | 5,9% | P972P (CCC→CCA) | *J9265_03690 →* | hypothetical protein |
| Chr 1 | 909 | C→T | 5,5% | V280V (GTC→GTT) | *dnaA →* | Chromosomal replication initiator protein DnaA |
| Chr 1 | 1254570 | T→C | 5,5% | N52S (AAC→AGC) | *ydcV ←* | Inner membrane ABC transporter permease protein YdcV |
| Chr 1 | 1934332 | G→A | 5,3% | W59* (TGG→TGA) | *J9265_08690 →* | hypothetical protein |
| Chr 1 | 1453032 | A→G | 5,2% | L82P (CTC→CCC) | *syrM1 ←* | HTH‑type transcriptional regulator SyrM 1 |
| Chr 2 | 750227 | C→A | 5,2% | H359N (CAT→AAT) | *rnb →* | Exoribonuclease 2 |

a : Gene or locus name corresponding to *Vibrio cholerae* A1552 chromosome 1 [CP072847.1](https://www.ncbi.nlm.nih.gov/nuccore/CP072847.1) or chromosome 2 : [CP072848.1](https://www.ncbi.nlm.nih.gov/nuccore/CP072848.1).

**Table S4 Mutation accumulated in MO10 V2**

| **Chromosome** | **Position** | **Mutation** | **Frequency** | **Annotation** | **Gene name or locus** | **Description** |
| --- | --- | --- | --- | --- | --- | --- |
| Chr 1 | 375972 | C→T | 100% | Q173* (CAA→TAA) | *mlaF →* | Intermembrane phospholipid transport system ATP‑binding protein MlaF |
| Chr 1 | 2467074 | Δ11 bp | 100% | coding (1013‑1023/1446 nt) | *dacB →* | D‑alanyl‑D‑alanine carboxypeptidase DacB |
| Chr 1 | 1774887 | T→G | 22,7% | L57* (TTA→TGA) | *hutI →* | Imidazolonepropionase |
| Chr 2 | 634938 | G→T | 6,5% | L176L (CTC→CTA) | *fbpC ←* | Fe(3+) ions import ATP‑binding protein FbpC |
| Chr 1 | 783279 | C→A | 6,0% | A363D (GCT→GAT) | *flaD_2 →* | Flagellin D |
| Chr 1 | 2999226 | G→T | 6,0% | Q48K (CAA→AAA) | *ilvC ←* | Ketol‑acid reductoisomerase (NADP(+)) |
| Chr 1 | 2285167 | C→T | 5,9% | R238* (CGA→TGA) | *rdgC →* | Recombination‑associated protein RdgC |
| Chr 2 | 1031122 | C→G | 5,8% | W399C (TGG→TGC) | *menE_2 ←* | 2‑succinylbenzoate‑‑CoA ligase |
| Chr 1 | 564552 | C→A | 5,6% | L341I (CTT→ATT) | *thrC →* | Threonine synthase |
| Chr 2 | 569937 | G→T | 5,5% | S17* (TCG→TAG) | *sdaA_2 ←* | L‑serine dehydratase 1 |
| Chr 1 | 1245901 | G→T | 5,4% | A80E (GCG→GAG) | *ynjF ←* | Inner membrane protein YnjF |
| Chr 1 | 2273204 | G→T | 0,052 | R107R (CGC→CGA) | *pstB ←* | Phosphate import ATP‑binding protein PstB |
| Chr 1 | 2454795 | C→A | 5,2% | E221* (GAG→TAG) | *infB ←* | Translation initiation factor IF‑2 |
| Chr 1 | 2627880 | G→T | 5,2% | L137L (CTC→CTA) | *argP_1 ←* | HTH‑type transcriptional regulator ArgP |
| Chr 2 | 695723 | G→T | 5,2% | W113C (TGG→TGT) | *KAF59_17225 →* | hypothetical protein |
| Chr 1 | 1178677 | G→T | 5,0% | M51I (ATG→ATT) | *glgC_1 →* | Glucose‑1‑phosphate adenylyltransferase |

a : Gene or genome locus corresponding to *Vibrio cholerae* MO10 chromosome 1 : [CP072849.1](https://www.ncbi.nlm.nih.gov/nuccore/CP072849.1) or chromosome 2 : [CP072850.1](https://www.ncbi.nlm.nih.gov/nuccore/CP072850.1).

**Table S5 Mutation accumulated in MO10 V8**

| **Chromosome** | **Position** | **Mutation** | **Frequency** | **Annotation** | **Gene name or locus ^a^** | **Description** |
| --- | --- | --- | --- | --- | --- | --- |
| Chr 1 | 875871 | Δ371 bp | 100% |  | *ccmH-mlaA* | Cytochrome c-type biogenesis protein CcmH - Intermembrane phospholipid transport system lipoprotein MlaA |
| Chr 2 | 644437 | A→G | 20,8% | A64A (GCT→GCC) | *KAF59_16975 ←* | hypothetical protein |
| Chr 1 | 2448239 | G→T | 16,1% | Q22K (CAG→AAG) | *nlpI ←* | Lipoprotein NlpI |
| Chr 1 | 183145 | G→T | 6,2% | E45* (GAG→TAG) | *epsL →* | Type II secretion system protein L |
| Chr 1 | 838058 | G→T | 6,2% | E296* (GAG→TAG) | *sucC →* | Succinate‑‑CoA ligase [ADP‑forming] subunit beta |
| Chr 1 | 1848806 | T→C | 6,2% | T221A (ACT→GCT) | *hisF ←* | Imidazole glycerol phosphate synthase subunit HisF |
| Chr 1 | 1322424 | A→G | 6,1% | F52L (TTC→CTC) | *KAF59_06085 ←* | hypothetical protein |
| Chr 1 | 2539488 | G→T | 5,5% | E90* (GAG→TAG) | *yhcB →* | Inner membrane protein YhcB |
| Chr 1 | 1885682 | C→A | 5,2% | A28S (GCC→TCC) | *KAF59_08560 ←* | hypothetical protein |

a : Gene or genome locus corresponding to *Vibrio cholerae* MO10 chromosome 1 : [CP072849.1](https://www.ncbi.nlm.nih.gov/nuccore/CP072849.1) or chromosome 2 : [CP072850.1](https://www.ncbi.nlm.nih.gov/nuccore/CP072850.1).

```
>ccmH-vacJ MO10-V8
 Length of sequence-      1661
 Threshold for promoters -  0.20
 Number of predicted promoters -      4
 Promoter Pos:    959 LDF-  3.51
 -10 box at pos.    944 TTTCACACT Score    34
 -35 box at pos.    923 TTAAGG    Score    14
 Promoter Pos:   1554 LDF-  1.97
 -10 box at pos.   1539 CCTTACAGT Score    35
 -35 box at pos.   1521 TTACAA    Score    32
 Promoter Pos:    598 LDF-  1.60
 -10 box at pos.    583 CTGGATAAT Score    47
 -35 box at pos.    565 ATGAAG    Score    24
 Promoter Pos:    268 LDF-  0.91
 -10 box at pos.    253 CAGGAAAAT Score    33
 -35 box at pos.    230 TTCCAA    Score    35

 Oligonucleotides from known TF binding sites:

 For promoter at    959:
        crp:  TCACACTT at position     946 Score -  11
       carP:  CACTTTTT at position     949 Score -   8
       lexA:  TTTTTTTA at position     952 Score -  16
 For promoter at   1554:
      phoB3:  TCCTTACA at position    1538 Score -  13
 No such sites for promoter at    598
 No such sites for promoter at    268
```

Figure S1. BPROM prediction for MO10-V8 *mlaA** [36]. In MO10-V8 mutation in *ccmH* and *mlaF* resulted in apparition of a new ORF at position +624 *see Supplemental material and methods*.

```
> mlaF*_MO10-V2
 Length of sequence-       804
 Threshold for promoters -  0.20
 Number of predicted promoters -      2
 Promoter Pos:    108 LDF-  3.56
 -10 box at pos.     93 aggtaaagt Score    58
 -35 box at pos.     68 atgaca    Score    36
 Promoter Pos:    508 LDF-  3.25
 -10 box at pos.    493 atgtatgat Score    66
 -35 box at pos.    470 ttgcac    Score    33

 Oligonucleotides from known TF binding sites:

 For promoter at    108:
        ihf:  CTTCGGGA at position     119 Score -  12
 For promoter at    508:
     rpoD18:  TGTATGAT at position     494 Score -   7
```

**Figure S****2. BPROM prediction for MO10-V2 *mlaF** promoters [36].** In MO10-V2 mutation in *mlaF* resulted in apparition of a new ORF at position +532 *see Supplemental material and methods*.

```
>dacB* A1552-V6
 Length of sequence-      1434
 Threshold for promoters -  0.20
 Number of predicted promoters -      3
 Promoter Pos:    306 LDF-  2.44
 -10 box at pos.    291 ttggatcat Score    33
 -35 box at pos.    270 tttaat    Score    36
 Promoter Pos:   1141 LDF-  2.19
 -10 box at pos.   1126 tggtttaat Score    56
 -35 box at pos.   1105 ttatca    Score    30
 Promoter Pos:    728 LDF-  1.59
 -10 box at pos.    713 cactaaaat Score    63
 -35 box at pos.    689 tagccc    Score     2

 Oligonucleotides from known TF binding sites:

 For promoter at    306:
     rpoD17:  GACCCCAC at position     313 Score -  10
 No such sites for promoter at   1141
 No such sites for promoter at    728
```

**Figure S3. BPROM prediction for A1552-V6 *dacB** promoters [36].** In A1552-V6 mutation in *bacA* resulted in apparition of a new ORF at position +1067 *see Supplemental material and methods*.

**Supplemental Material and methods**

A1552-V2, MO10-V2 and MO10-V8 mutations

Full open reading frame sequences of *ihfA*, *dacB*, *ccmH-vacJ* and *mlaF*. Sequences starts at first nucleotide of indicated gene ORF.

> *ihfA** A1552-V6

ATGGCGCTCACAAAGGCCGAATTGGCTGAAGCCCTGTTCGAACAGCTCGGCATGAGCAAGCGGGATGCCAAGGATACGGTTGAGGTGTTTTTTGAAGAAATTCGTAAAGCACTCGAAAGTGGCGAACAGGTAAAACTCTCCGGTTTTGGTAATTTTGACCTACGAGATAAAAATGAACGTCCGGGTCGAAACCCTAATATTCCTATTACCGCTCGACGTGTCGTAACGTTCCGCCCAGGGCAAAAATTGAAAGCCCGTGTCGAGAACATCAAAGTCGAAAAATAA

> *dacB** A1552-V6

ATGCTTTTTCGCTTCATACCTGTTTGGTTACTCTCTATTAGCGGTTTTCTAATAGCCTCTCCGATTTACGCACAAACACCATTGACTGCCGCAACGACTAAACTTCCTCAAGGGGCACGTTATAGCCTATTGATTGAAGATGTCGCATTACAGCAGAACACTCTCGAACTCAATACTCATCTGTACTATCCCCCCGCTAGCACCCAGAAGATTTTGACGGCACTCGCCGCAAAATTAGAACTGGGTGATAAGTTTCGCTTTCACACTGATTTAATGCGTTCAGGGCAAGATTGGATCATTCGCTTTTCAGGCGACCCCACCCTGACCACCGCAGATTTAACGACATTGCTCAAAGCGATGAAAGCGCAAAGTGGCGGTAGGATTGAGGGCGATTTGTGGCTGGATAATAGTGTATTTAGTGGATATGAGCGTGCGGTAGGCTGGCCATGGGATATTTTAGGCGTCTGCTATAGCGCCCCAGCCAGTGCCATCAACCTCGATGCTAACTGTATCCAGGCGTCTATTTATACCGAACCACAAGGTAAAACGCGCGTTTATGTACCAGAGCACTACCCTGTGCATGTTCAGTCGCAAGCCATCAGCGTGACACAGAGCGAACAAGAGAGTTTACTGTGTGACTTAGAACTGACGGCAACGCCTGAGAATCACTATACGCTGGATGGCTGTTTAGCCCTACGAGACAAACCCTTACCACTAAAATTTGCGGTGCAAGATACTGGGATCTACACCCAGCGAGTGGTCTATCGTCTCCTCAGCCAGCTAAACATCGAGCTCAAAGGGAAGATAAAAGTCGGTAAAGCAAATACCAAACAAGCGCAGAAAATCGCTTCTCATCACTCCCAGCCGCTGCCTGTGTTACTGAAAACCATGTTGCAAGAGTCCGACAACCTGATCGCCGATACCTTGACCAAAGCCTTGGGACACCGTTTTTACTCTCAACCCGGTAGCTTTACCAACGGAACACAAGCCATTAAACAGATTTTTTACTCGCTCATTAGAAGATACTCAGCTCGCCGATGGCTCTGGCCTTTCACGTAATAACCGGATGCGTCCACAAGTGATGCTGGAAACTCTTCGCTACCTTTATCAGCACGAAGCTGAGCTTGGTTTAATTGCTATGCTGCCTTCAGCGGGAGAATCGGGCACTTTGCAATATCGACGCAGTATGCGTGCGCCGCAAATCAGTGGCCAAATTAAAGCGAAAAGTGGTTCACTTTATGGCACTTACAATATGGCGGGCTTTGTGATGGACGAAAATCAGCGCCCTAAGACTCTGTTTGTTCAATTCGTCACCGACTATTTCCCTCCGAGATCCAATCCTGAGGTAGCGGTTGAGCCGCCGATTATCCAGTTTGAAACTCAGCTCTATCAAGAGCTTATTCAGTTTAATCGTTTGGCATCTAAGCCAAACTA

>*ccmH**-*vacJ** MO10-V8

ATGTGGATGTTTTGGATCTCGACCCTATTACTGGTGGCGATTGCGGTGGTTTTCGTCATCATTCCGTTTATTCAAAAGCGTGCGAATAACGATCAGGCTTTGCGCGATGAGCTGAATAAAGCGTTTTACAAAGACCGCTTGAAAGAGCTTGAAGAGGAAACCGAAGAAGGCATTGTTGCCGATCAACAAGATTTGATTGCCGACTTAAAACAGACTCTGCTTGACGACATTCCAACCCAGCAAAAACATCAGCAGGAAAATCGTGTTTCACTGTGGATGGTTGCCCTGCCTTCAGTATTGTTGGTAGTCGGATTGAGTTATGCGCTGTACGCCAAGTTTGGTCACTATCAGCATGTTCAGGCTTGGCAGCAAGTGTCAGCACAACTGCCTGAATTGTCAAAAAAACTCATGTCGCCACAAGCGGAACTCAGTGACGAAGAGATGAATGATTTGACGTTGGCACTGCGCACTCGACTGCATTATCAGCCTGATGATGTTACCGGTTGGTTGTTGCTGGGTCGGATTGGCCTTGCTAATCGCGATCTGGAAACCGCGATTGGCGCGATGAAGAAAGCTTTTGCTCTGGATAATGAAGATCCGGATGTGAAATTTGGTTACGCACAAGCTTTGATGCTTTCGAATGATCCTGTCGACCAGCAAGAAGCGAAGTCGATTCTGCTCAAGTTAGCCCAACGTGGTTATGCTGATTTACGCGTCTATTCATTATTGGCGTTTGATGCTTTTGAAAGTGGAGATTTTCCTGCTGCAATCAAGTACTGGAGTTTGATGCAACAAGCGATTGGTCCTGACGATGCTCGTTATGAGATGCTCAGCCGCAGTATTGAAAGCGCTCGTAAGAGAATGGGCGAGGGCATGGCAGAGGGTCAATCGGTGAAAGTCACCATTAATCTAGGCGAGCAGGTTAAGGTTGATCCTAACGCAGTTTCACACTTTTTTTAGTTGGGTGCAGCAGTGCACCTGATGACTCCTCCCCTCATTCGCAGGTGAACGATCCTCTGGAAAGTTTCAACCGGCAAATGTGGACAATTAACTATGACTACCTAGACCCTTATGTGGTGCGTCCGGTCTCTCTATTTTATGTCGGTTATGTACCTAAGCCTGTACGCAGTGGCATTGCCAACTTCCTCTCTAACTTAGACGAGCCTGCCAGCATGGTGAATAACCTGCTGATGGGCAATGGGACAAAAGCGGTCGATCACTTTAATCGTTTTTGGATTAATACCAGCTTTGGTTTACTCGGTTTGATTGATATCGCTTCTGAAGCAGGCATCAAAAAATACGATGATAAGGCGTTCAGTGATGCGGTAGGCCATTACGGTGTGGGCAATGGCCCCTATTTAATGGTCCCAGGTTATGGTCCCTATACGGTACGCGAAGTGACCGATGTGGTGGATGGCATGTATTTCCCGCTTGCCTATCTCAATATCTGGGCTGGGGTCGGCAAATGGGCACTTGAAGGCATGGAAACGCGCGCTGCGTTAGTTTCGCAAGAGGCCTTATTACAAGACTCACCTGATCCTTACAGTTTGGCTCGCGATGCTTATCTCCAACGGCAAGCTTTCAAAGCGGAGATCCAAGTGGATGACTATGACCCTGAGGAAGAAGAGTATCTCGATGAGTATTTAAATGAAGGGTTATGA

>*mlaF** MO10-V2

ATGTCTCAATCTGACTTAGTCACCATCAAAAATTTGCGTTTTTCGCGCTCGCAGCGCGTCATTTTTGATGACATAGATCTGCATGTCCCCAAAGGTAAAGTGACAGCAATTATGGGGCCTTCGGGAATCGGTAAAACCACACTGCTGCGTTTGATCGGCGGTCAACTCCTGCCAGAACAGGGAGAGATCTGGTTCGATGGTGAAAATATTCCCACCCTCAGTCGCCGCAAACTGTATCGTGCTCGTAAGAAGATGAGCATGCTGTTCCAATCAGGCGCGCTGTTTACCGATCTTAATGTGTTTGACAATGTGGCTTACCCATTGCGCGAGCATACCGAACTTGATGAAGCCATGATTAAAACCTTGGTGCTGCTGAAATTAGAGGCGGTTGGACTGCGTGGTGCTGCGTATTTAATGCCTAGTGAGCTTTCAGGCGGTATGGCGCGCCGCGCCGCACTGGCAAGGGCTATTGCACTCGATCCTGAGCTCATCATGTATGATGAGCCGTTTGTCGGATAAGATCCGATCACCATGGGTGTACTGGTTGAACTGATCCGTAACCTTAATCGAGCCTTGGGTGTCACCTCTGTGGTGGTATCGCACGATGTACCGGAAGTGATGAGCATTGCGGATTGGGTTTATCTGTTGGCCGATGGTAAGGTGATTGCGCAAGGTTCACCTCAAGCATTGCGCGACAACCCTGATCCGCGTGTACAACAATTTTTATGCGGCGATGCAGATGGCCCTGTGCCATTTCGTTTTCCTGCGCAGCCGATAGAACAGGAGCTGTTTAGTGCTAAATGA
